# Novel feeding and mating behaviors of a population of nautiluses, *Nautilus belauensis*, in Palau

**DOI:** 10.1101/622456

**Authors:** Gregory J. Barord, Rebecca L. Swanson, Peter D. Ward

## Abstract

The nautiloid lineage extends back nearly 500 million years but today, is represented by only two living genera, *Nautilus* and *Allonautilus*. Behavioral observations of these living nautiluses have improved our understanding of how nautiloids, and ammonoids, behaved and interacted in their environment. These behaviors may also help to inform conservation practices. Here, we describe feeding and mating behaviors in wild nautiluses not reported from any other population. In Palau, *Nautilus belauensis* was observed actively preying on a large, living crab (*Chaceon* sp.) and performing courtship-like behaviors prior to mating. These behaviors occurred across multiple nights and from different nautiluses, suggesting that the behaviors are characteristic of at least a subset of the population, if not the entire population. Perhaps the behaviors exhibited by the Palauan nautiluses are an outlier and simply a localized characteristic of a far-removed population. Or, perhaps these apparent abnormal behaviors of Palauan nautiluses are what all nautiluses across the Indo-Pacific should be exhibiting. If the latter explanation is correct, we can start to address the potential causes of the behavioral differences, such as population size, habitat type, and prey availability. In either case, this apparent behavioral plasticity may have also been a reason that the nautiloid lineage has been able to survive throughout millions of years of environmental changes. Today, these behavioral observations could prove to be a valuable conservation tool to protect species and environments, especially in the deep-sea ecosystem the nautiluses inhabit.

## Introduction

Living nautiluses (extant species of *Nautilus* and *Allonautilus*) are the last vestiges of an ancient 500-million-year-old nautiloid lineage that have provided invaluable information about morphologically similar, but extinct, taxa^1,2^. Behaviors of living nautiluses have informed how ancient ecosystems may have functioned, but as nautilus populations continue to decline^3,4,5^, these behaviors may be more informative of healthy and unhealthy populations. Related to octopuses, squid, and cuttlefish, living nautiluses most closely resemble their ancient nautiloid counterparts, characterized by a hard, external shell. Today, nautiluses inhabit the deep-sea coral reef slopes across the Indo-Pacific from 0-700 meters^6^. Their habitat is constrained by warm, surface water temperatures of 25 °C, depth implosion limits at 800 meters, and a nektobenthic lifestyle^7,8,9^. Nautiluses locate food in the deep-sea using their large olfactory organs and dozens of sticky tentacles^10^ to cue in on dead prey items. Unlike many cephalopods, nautiluses are iteroparous, mature late, produce few offspring, and the embryos have long developmental times^11,12,13,14^. Nautiluses are highly prized for their ornamental shell that can be sold whole, or made into jewellery, furniture inlay, or other curio items^4^, significantly depleting populations^5,15^. To survey nautilus populations, baited remote underwater video systems (BRUVS) have become the standard for use with nautiluses and their deep-sea habitat, or “nautilus zone”, between 100-700 meters. In addition to providing population data, the BRUVS also record behaviors that otherwise would not be known and in Palau, the video data showed nautilus behaviors not reported from any other populations.

The nautiluses of Palau represent the northernmost range of Family Nautilidae with no known historical fisheries in the region. In the literature, these nautiluses have been described morphologically as *Nautilus belauensis*^16^ and recent genetic analyses have provided increased support for *N. belauensis* as a distinct species^17,18^, apart from the widespread *Nautilus pompilius*, though this has not been settled. However, for this paper, we will refer to the nautiluses of Palau as *Nautilus belauensis* to avoid confusion between other populations of nautiluses. Belauensis is also commonly known within practice and the culture of Palau (or Belau) to describe the nautiluses therein.

Nautiluses are highly adapted to locating food in the deep-sea. They have large olfactory organs, called the rhinophores^19^, and dozens of protractible and retractable tentacles^10^. Nautiluses have been described as predators, scavengers, and opportunistic predators/scavengers^20,21^ but no direct behavioral evidence is available on their actual diet in the wild. In aquariums, nautiluses are fed a varied diet centred on prepared, frozen shrimp^22^. Nautiluses are commonly displayed with other marine organisms, though there are no published reports of active predation on these co-habitants. Using time lapse photography, nautiluses in Palau did not prey on live prey items, but traps with shrimps, sharks, and fishes did show nautilus bite marks^21^, though it is not clear what condition the organisms were in, dead, near-dead, or alive, or if the bites were predatory.

## Materials and Methods

### Baited Remote Underwater Video Systems

Baited remote underwater video systems (BRUVS) were used record the behavior of nautiluses, and other species, attracted to a bait source over a fixed period. Each baited remote underwater video system, or BRUVS, is composed of one HD-camcorder in an underwater housing and one LED light source in an underwater housing affixed to a steel frame (1m^3^) with a bait stick extending off the front 2 meters and a rope attached to a surface buoy. The BRUVS were deployed at dusk to a depth of 250-300 meters using a Furuno Fish Finds (MODEL FCV587) and retrieved 12 hours later at dawn. Upon retrieval, the video data was transferred to an external hard drive and the BRUVS were fitted for deployment later that day. The video data was later analysed for nautilus behaviors, habitat type, species composition, and population abundance. All methods were carried out in accordance with relevant guidelines and regulations. No nautiluses, or other animals, were removed from their habitats or handled in any way with only the video data taken.

## Results and Discussion

### Feeding Behaviors

Previous observations from BRUVS in all other populations of nautiluses surveyed show a negative, or at least passive, response to potential living prey items^23,24^. In each case, the nautiluses showed no positive response to live shrimp, crabs, or fishes near the bait source, and seemingly within the grasp of the nautiluses’ tentacles (Figure 1). In Palau, a *N. belauensis* was observed performing predatory-like behaviors on a large crab, *Chaceon* sp. (Fig 2a-d). The nautilus jets to the posterior of the crab (Fig 2a), attaches with its tentacles (Fig 2b), attempts to jet away with the crab (Fig 2c), and finally detaches after the crab attempts to pinch the nautilus (Fig 2d). Thereafter, the nautilus continues to “hunt” the crab until both are out of frame. On a separate night, multiple nautiluses were recorded performing similar predator type behaviors on the same crab species.

**Figure 1.**
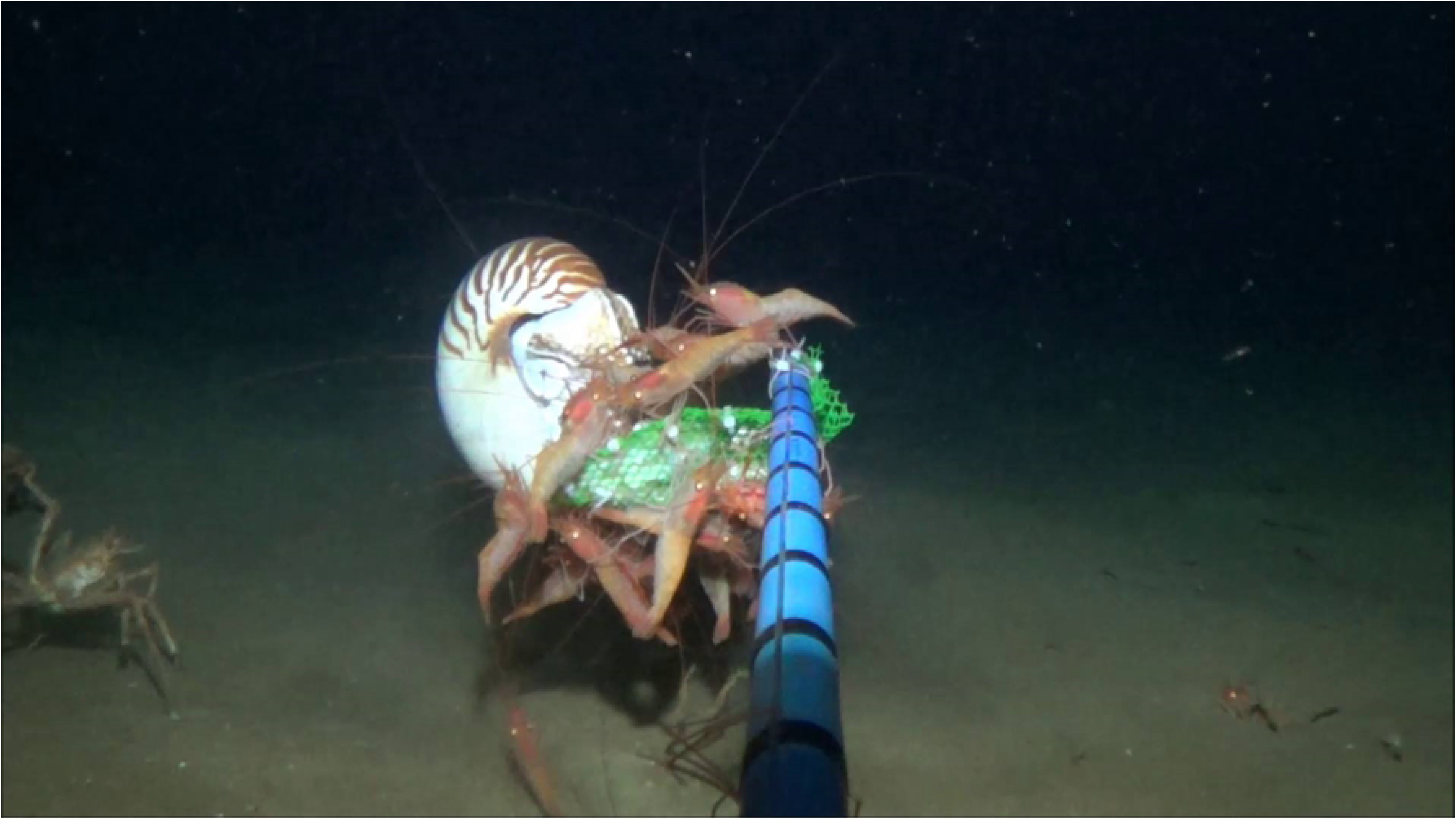
*Nautilus pompilius* feeding on raw chicken bait at 300 meters in the Philippines.

**Figure 2.**
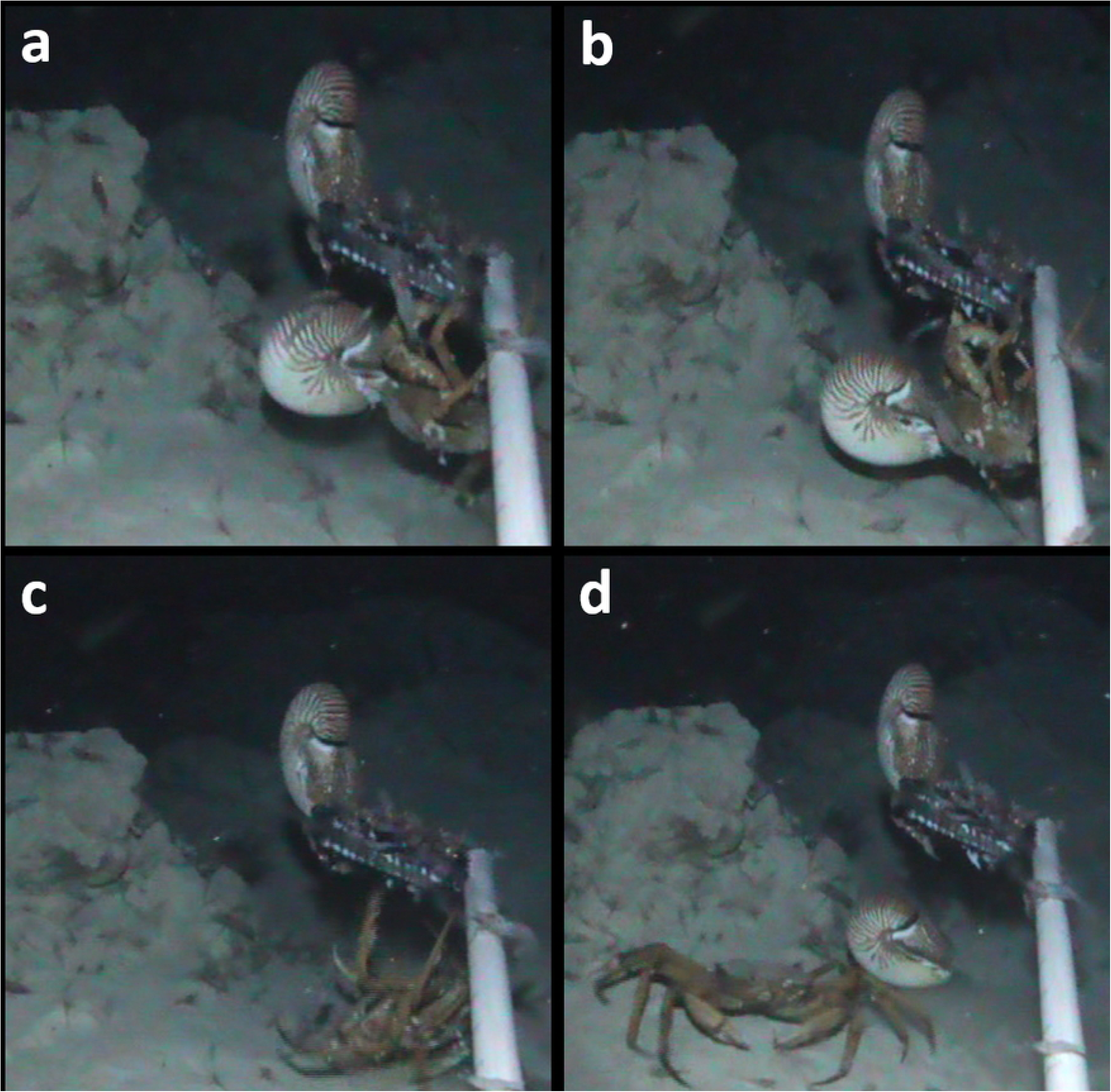
(a-d). *Nautilus belauensis* in Palau displaying predatory behaviors on a large crab, *Chaceon*, by latching on with its tentacles (2a), attempting to jet away (2b), being pinched by the crab (2c), and moving into the defensive position with tentacles contracted in shell (2d).

Aside from predation on the crab, the behaviors could be interpreted differently. First, the predation may not be on the actual crab, but parasites that are living on the crabs’ carapace. There are many examples of marine fishes and crustaceans that groom larger fishes of parasites in a symbiotic relationship^25,26^. However, the behaviors of the nautiluses and crabs do not support any type of evolved symbiosis for parasite removal; the nautilus attempts to jet away with the crab while the crab eventually pinches the nautilus with its chelipeds. An additional explanation could be that the nautilus was attempting to lay one of its few eggs on a moving, hard substrate, which was a crab in this case. In aquariums, nautiluses have laid eggs on other nautiluses on rare occasions (Pers Comm). But again, the initial behavior of latching on to the crab and attempting to jet away is more characteristic of finding food and taking it somewhere else to consume, rather than egg-laying. Finally, these behaviors may be related to competition-related behaviors between the nautilus and crab as both are scavenging on the bait, the nautilus behavior may be defensive as to protect its food source.

### Mating Behaviors

A second, previously undescribed behavior was also recorded from the BRUVS. Nautilus reproduction has most commonly been studied in laboratory and aquarium settings because of the limited access to the deep-sea, nautilus zone. Recently, a growing number of wild observations utilizing BRUVS are becoming available^3,5,24^. Based on our current understanding of nautilus reproduction, mating in nautiluses can best be described as a “trial and error” strategy with males attempting to mate with any sex. As nautiluses approach the bait source, they usually will feed first, and then locate a suitable mate. Male nautiluses are attracted to both male and female nautiluses, whereas female nautiluses are attracted to males, but repelled by other females^27^. These behaviors are present in aquarium observations with known sexes and appear in wild video observations. If we apply this proxy to wild populations, it is possible to discern potential males from the video. *Nautilus belauensis* shows similar mating behaviors to other populations throughout each video, across sampling days, and across individuals. However, the difference in *N. belauensis* is that the presumed male nautilus curls its tentacles along the posterior of the shell of each nautilus it encounters (Fig 3), a behavior not recorded from any other population of nautiluses across the Indo-Pacific.

**Figure 3.**
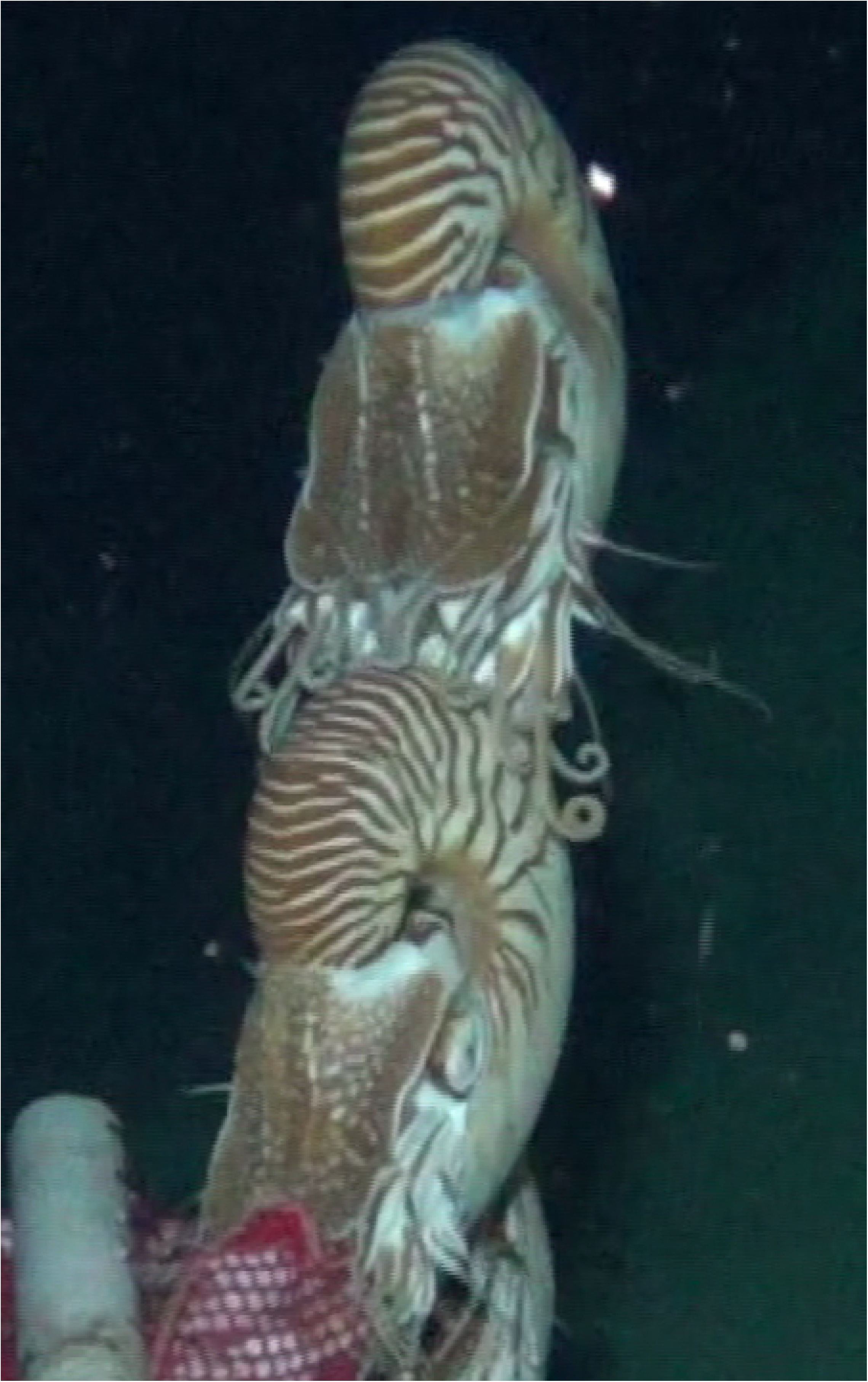
*Nautilus belauensis* performing courtship-like behaviors recorded from BRUVS typified by the curling of the tentacles tips on the shell of other nautiluses.

This curling behavior could simply be an artefact of tentacle movements, characteristic of the Palauan nautiluses and have no importance in mate selection and reproduction. If this were the case, we would expect to also observe the curling tentacles during feeding behaviors and other movements, which we do not. We cannot determine what the curling of tentacles on the nautilus shell is signalling from these observations and whether the nautilus with curled tentacles is sending a signal, or vice versa. What we can say is that mating (Figure 4) does occur after the tentacle curling behaviors of the nautiluses.

**Figure 4.**
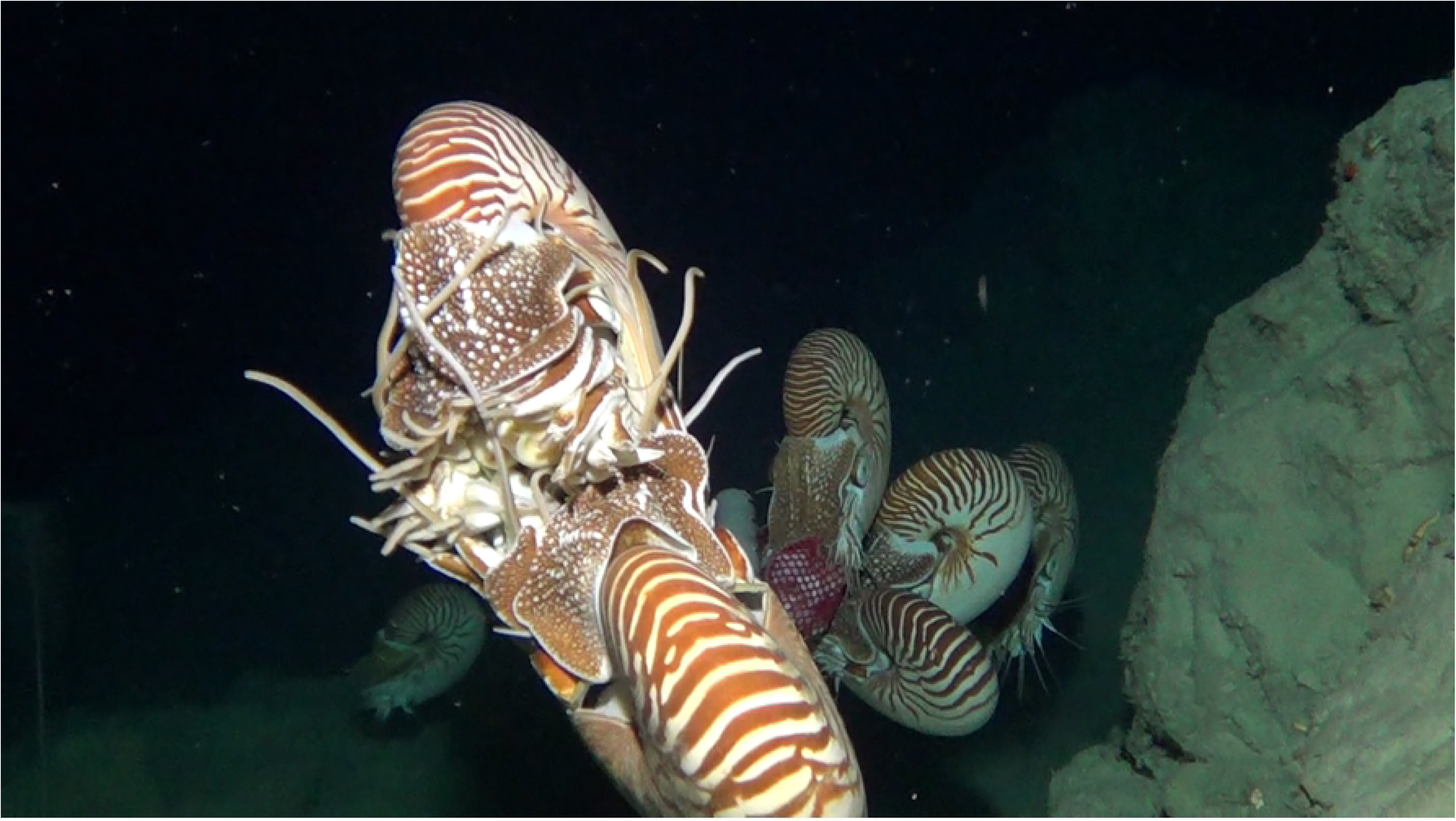
*Nautilus belauensis* mating, tentacles to tentacles, after displaying the courtship-like behaviors of curled tentacles on the shell.

Particularly related to behavior, we understand that the BRUVS rely on a fixed bait source to attract nautiluses, and other organisms, that potentially would not meet otherwise. However, this practice may mimic precisely how scavengers locate food in the deep-sea, where food is scarce. Deep sea scavengers may rely on windfall events when large prey items die and drift to the bottom of the sea^28^. The increased number of scavengers may, in turn, attract larger predators, different species, and greater abundances of specific species. If this is the case, then we can draw more concrete conclusions about the deep-sea ecosystem and relationships of animals utilizing BRUVS.

Taking these two behaviors together, we can assume that these are isolated anomalies of *N. belauensis* that evolved independent of other populations. Or, we can conclude that these behaviors are, in fact, indicators of a normal, healthy population of nautiluses suggesting a complex behavioral plasticity of nautiluses under different environmental conditions. In Palau, there are no commercial fishing activities of nautiluses. Their catch rates and population abundance (GJB, RS, & PDW, In Prep) are greater than other populations and may not just signify a healthy nautilus population, but a healthy nautilus zone, between 100-700 meters. Increased populations of nautiluses would need to be supported by increased numbers of prey items and resources. With additional resources, nautiluses may be able to shift to become more active predators, expending more energy to locate higher valued prey items with a greater energy return. Increased resources may also impact nautilus behavior by bringing nautiluses, which are presumed to be solitary, in closer proximity to other nautiluses during feeding events. The proximity to other nautiluses, then, may provide for a type of sexual selection, either by the male or female. Sexual selection has been described in other cephalopods, including *Sepia apama* where the females choose their mates^29^. In contrast, populations with fewer nautiluses may rely more heavily on a strict scavenging strategy to conserve energy while searching out prey items and when encountering other nautiluses, forego any courtship behaviors to instead, simply pass on their genes and mate.

## Conclusions

Looking to the past, the behaviors of *N. belauensis* could help inform our understanding of how the nautiloid ancestors may have behaved and interacted with their ecosystem. Few species can truly serve as a model for how its fossilized ancestors may have behaved, while also serving as a sign of what the future may be. Utilizing behavior as a conservation tool to assess population and ecosystem health is useful, particularly in areas like the nautilus zone that are difficult to survey scientifically, but relatively easy to exploit by fishermen.

## Acknowledgments

The authors thank the Coral Reef Research Foundation for assistance on equipment construction and gear deployment.

## References

1. Saunders, B. & Landman, N. Nautilus: The Biology and Paleobiology of a Living Fossil. Reprint with additions. New York, Plenum Press, doi:10.1007/978-90-481-3299-7 (2010).

2. Ward, P. The Natural History of Nautilus. Winchester, Allen and Unwin, 281 p (1987).

3. Dunstan, A., Bradshaw, C., & Marshall J. Nautilus at risk-estimating population size and demography of *Nautilus pompilius*. PLoS ONE 6, e16716 (2011).

4. DeAngelis, P. Assessing the impact of international trade on chambered nautilus. Geobios 45, 5–11 (2012).

5. Barord, G.J., et al. Comparative population assessments of *Nautilus* sp. in the Philippines, Australia, Fiji, and American Samoa using baited remote underwater video systems. PLoS ONE 9(6), e100799 doi:10.1371/journal.pone.0100799 (2014).

6. Dunstan, D. & Ward, P. Vertical distribution and migration patterns of *Nautilus pompilius*. PLoS ONE 6, e16311 (2011).

7. Ward, P.D., Greenwald, L., & Rougerie. Shell implosion depth for living *Nautilus macromphalus* and shell strength of extinct cephalopods. Lethaia, 13-18 (1980).

8. Carlson, B.A. Collection and aquarium maintenance of *Nautilus*. In: Saunders, W.B. & Landman, N.H. (eds), Nautilus: The Biology and Paleobiology of a Living Fossil. New York, Plenum Press, 563–578 (2010).

9. Ward, P.D., & Martin, A.W. Depth distribution of *N. pompilius* in Fiji and *N. macromphalus* in New Caledonia. Veliger 22, 259–264 (1980).

10. Bidder, A.M. Use of tentacles, swimming, and buoyancy control in the pearly *Nautilus*. Nature 196, 451–454 (1962).

11. Landman, N.H., Cochran, J.K., & Saunders, W.B. Growth and longevity of *Nautilus*. In: Saunders, W.B., & Landman, N.H. (eds.), Nautilus: Biology and Paleobiology of a Living Fossil. New York: Plenum Press. pp. 401–420 (2010).

12. Saunders, W.B. Natural rates of growth and longevity of *Nautilus belauensis*. Paleobiology 9, 280–288 (1983).

13. Saunders, W.B. *Nautilus* growth and longevity: Evidence from marked and recaptured animals. Science 224, 990–992 (1984).

14. Okubo, S., Tsujii, T., Watabe, N., & Williams, D.F. Hatching of *Nautilus-belauensis* in captivity - culture, growth and stable-isotope compositions of shells, and histology and immunohistochemistry of the mantle epithelium of the juveniles. Veliger 38, 192–202 (1995).

15. Dunstan, A., Alanis, O., & Marshall, J. *Nautilus pompilius* fishing and population decline in the Philippines: A comparison with an unexploited Australian *Nautilus* population. Fish Res 106, 239–247 (2010).

16. Saunders, W.B. A new species of *Nautilus* from Palau. Veliger 24, 1–7 (1981).

17. Vandepas, L.E., Dooley, F.D., Barord, G.J., Swalla, B.J., & Ward, P.D. A revisited phylogeography of Nautilus pompilius. Ecology and Evolution 6:14, 4924-4935 (2016).

18. Combosch, D.J., Lerner, S., Ward, P.D., Landman, N.H., & Giribet, G. Genomic signatures of evolution in *Nautilus*—An endangered living fossil. Molecular Ecology 26:21 5923–5938 (2017).

19. Basil, J.A., et al. The function of the rhinophores and tentacles of *Nautilus pompilius* L. (Cephalopoda, Nautiloidea) in orientation to odor. Marine and Freshwater Behavior and Physiology 38:3, 209-221 (2005).

20. Ward, P.D. & Wickstein, M.K. Food sources and feeding behavior of *Nautilus macromphalus*. Veliger 23, 119–124 (1980).

21. Saunders, W.B. The role and status of *Nautilus* in its natural habitat: Evidence from deep- water remote camera photo sequences. Paleobiology 10, 469–486 (1984).

22. Barord, G.J. & Basil, J.A. Nautilus In: Inglesias, J., Fuentes, L., & Villanueva, R. Cephalopod Culture. New York, Springer Science 165–174 (2014).

23. Barord, G.J. On the biology, behavior, and conservation of the chambered nautilus, *Nautilus* sp. CUNY Academic Works, 96 p (2015).

24. Ward, P.D., Dooley, F., & Barord, G.J. *Nautilus*: biology, systematics, and Paleobiology as viewed from 2015. Swiss J Palaeontol 135, 169–185 (2016).

25. Johnson, P.T.J., et al. When parasites become prey: ecological and epidemiological significance of eating parasites. Trends in Ecology and Evolution 25, 362–371 (2010).

26. Cote, J.M. Evolution and ecology of cleaning symbioses in the sea. Oceanography and Marine Biology. 38, 311–355 (2000).

27. Basil, J.A., Lazenby, G.B., Nakanuku, L., & Hanlon, R.T. Female *Nautilus* are attracted to male conspecific odor. Bulletin of Marine Science 70, 217–225 (2002).

28. Smith, C.R. Food for the deep sea: utilization, dispersal, and flux of nekton falls at the Santa Catalina basin floor. Deep Sea Research Part A. Oceanographic Research Papers 32:4, 417-422 (1985).

29. Hall, K.C. & Hanlon, R.T. Principal features of the mating system of a large spawning aggregation of the giant Australian cuttlefish *Sepia apama* (Mollusca: Cephalopoda). Marine Biology 140, 533–545 (2002).

